# Efficient coding makes and breaks Weber’s law

**DOI:** 10.64898/2026.08.10.744043

**Authors:** Arthur Prat-Carrabin, Raphael Yamamoto, Samuel J. Gershman

## Abstract

Weber’s law is a rare quantitative regularity in psychology, yet its origins remain debated. Here we provide causal evidence that it arises from the more fundamental principle of efficient coding. This principle posits that representational resources are allocated according to stimulus frequencies: distributions skewed toward smaller stimuli thus result in discriminability decreasing with magnitude, as in Weber’s law. Skewing frequencies in the other direction—making large magnitudes more frequent than small ones—enabled us to invert this pattern, and to break Weber’s law. In discrimination tasks with three different sensory modalities, human subjects’ discriminability across stimuli was sensitive to the stimulus distribution, and this adaptation improved task performance. These findings establish efficient coding as a dynamic, organizing principle, explaining when and why Weber’s law holds.

---

Weber’s law states that the difference required to distinguish one stimulus from another (the just-noticeable difference, JND) is proportional to the magnitude of the stimulus^1^. Since its discovery more than 150 years ago, Weber’s law and its generalizations^1–3^ have successfully accounted for human discriminative ability with a very wide range of stimuli, including, for instance, sucrose concentration, acoustic frequency, and visual contrast^4–8^. Despite this enduring prevalence, the origin of Weber’s law remains debated. Most explanations point to the specific stochastic structure of sensory circuits^3,8–12^. But not all sensory modalities obey Weber’s law (in some cases the JND is a non-monotonic function of the stimulus magnitude^13,14^), and these mechanistic accounts do not explain *why* the neural code should be organized in such a way that discriminability decreases with magnitude. In short, Weber’s law is generally treated as a principle in and of itself. Here we demonstrate that a more fundamental principle, efficient coding, underlies Weber’s law, and we leverage this principle to experimentally suppress Weber’s law with three sensory modalities.

Efficient coding posits that neural coding resources should be allocated efficiently, and in particular as a function of the statistics of the encoded stimuli (the prior, *f* (*x*))^6,15–20^. This resource allocation determines the discriminability. Specifically, smaller JNDs are predicted for more frequent stimuli, a relation that efficient-coding models make quantitatively precise, e.g., JND 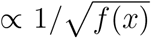. For many sensory variables, small magnitudes are more frequent than large magnitudes. Power laws (e.g., *f* (*x*) ∝ 1*/x*^2^), in particular, are ubiquitous in nature^21^. In these cases larger magnitudes should be allocated with less encoding resources, resulting in decreased discriminability, thus providing a candidate explanation for Weber’s law (JND ∝ *x*; see Supplementary Information for detailed derivations)^6,22–26^. Recent evidence suggests that efficient coding is not static, but is a dynamic property that adapts to the changing statistics of stimuli^27–29^. Therefore, if Weber’s law originates in the principle of efficient coding, then changing these statistics should alter the encoding of stimuli, which in turn should impact discrimination in a way that need not be consistent with Weber’s law.

We tested this prediction in three pre-registered experiments involving discrimination tasks with three different sensory modalities, where we manipulated the distribution of stimulus magnitudes. In particular, in some experimental conditions we made large magnitudes more frequent than small ones. This enabled us to break Weber’s law, suggesting that the law is not an immutable property of sensory systems. The changes we observed in participants’ discriminative behavior allowed them to enhance their performance (and reward) in the tasks, further supporting the proposal that sensory precision is modulated in order to favor better outcomes for the observer. Finally, the temporal dynamics of participants’ responses also suggested that the strength of sensory evidence for a magnitude increased with its relative frequency. Overall, our experimental results highlight the dynamic adaptability of sensory encoding, and establish efficient coding as the fundamental principle upon which Weber’s law depends.

## Results

We ran three discrimination tasks in which participants were asked to choose the greater of two stimuli of different length, brightness, or speed, presented to them in rapid suc-cession (Fig. 1A). In two experimental conditions, we manipulated the prior distribution from which the stimulus magnitudes were sampled: the *small-dominant* prior placed 80% probability on the lower half of the magnitude range, and 20% on the higher half; and conversely for the *large-dominant* prior (Fig. 1B). In each experiment, eight values of the difference between the two magnitudes, *x*_*right*_ *− x*_*left*_, were repeatedly tested, enabling us to quantify the participants’ discriminative power across the magnitude range (see Methods). Participants received a reward that increased with each correct response. To familiarize them with the relevant prior, the experiments started with 50 training trials in which feedback was provided, followed by 300 no-feedback test trials (on which all the analyzes below were based).

**Fig. 1:**
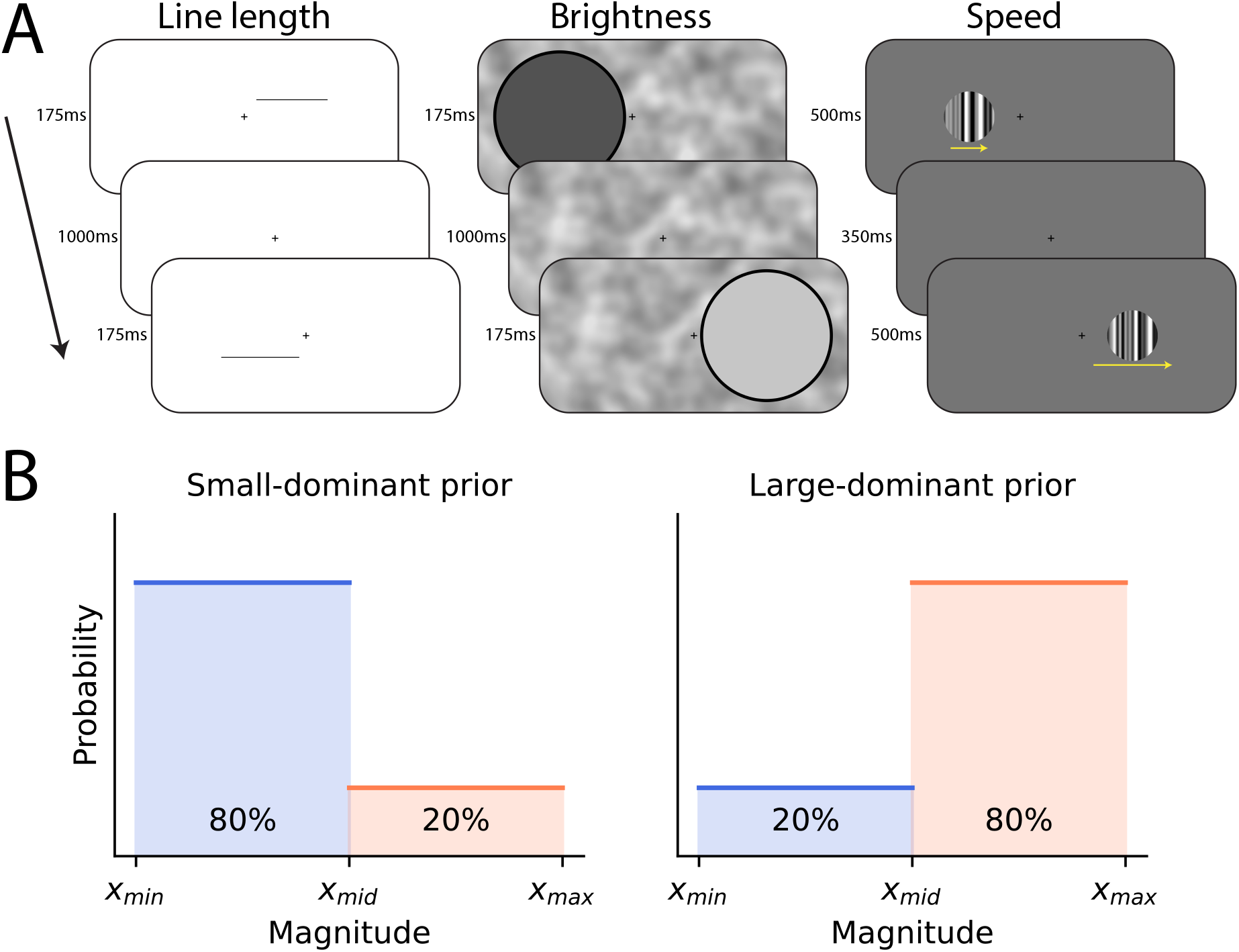
Discrimination tasks. **(A)** In three experiments, participants were asked which of two horizontal lines was longer, which of two patches of gray was brighter, and which of two horizontally-drifting gratings was moving faster. The presentations of the stimuli was sequential (separated by a blank screen showing only a fixation cross; total duration: 1350ms). Responses were self-paced. Yellow arrows (not shown in actual experiment) indicate drifting direction and speed. **(B)** Two experimental conditions differed by the distribution of presented sensory magnitudes (the prior). With the small-dominant prior, 80% of magnitudes were below the middle value of the magnitude range, and 20% above (left); with the large-dominant prior, 20% of magnitudes were below the middle value, and 80% above (right).

Larger differences in magnitudes yielded more correct choices (Fig. 2A-F). We split the trials depending on whether the average of the two stimulus magnitudes, 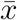 , was in the lower half of the range (*small-magnitude* trials) or the higher half (*large-magnitude* trials). With the small-dominant prior, the slope of the psychometric curve was steeper in small-magnitude trials than in large-magnitude trials, with significant differences for almost all distances *x*_*right*_ *− x*_*left*_ (Fig. 2A-C). In other words, consistent with Weber’s law, participants were less precise about larger magnitudes, with the small-dominant prior.

**Fig. 2:**
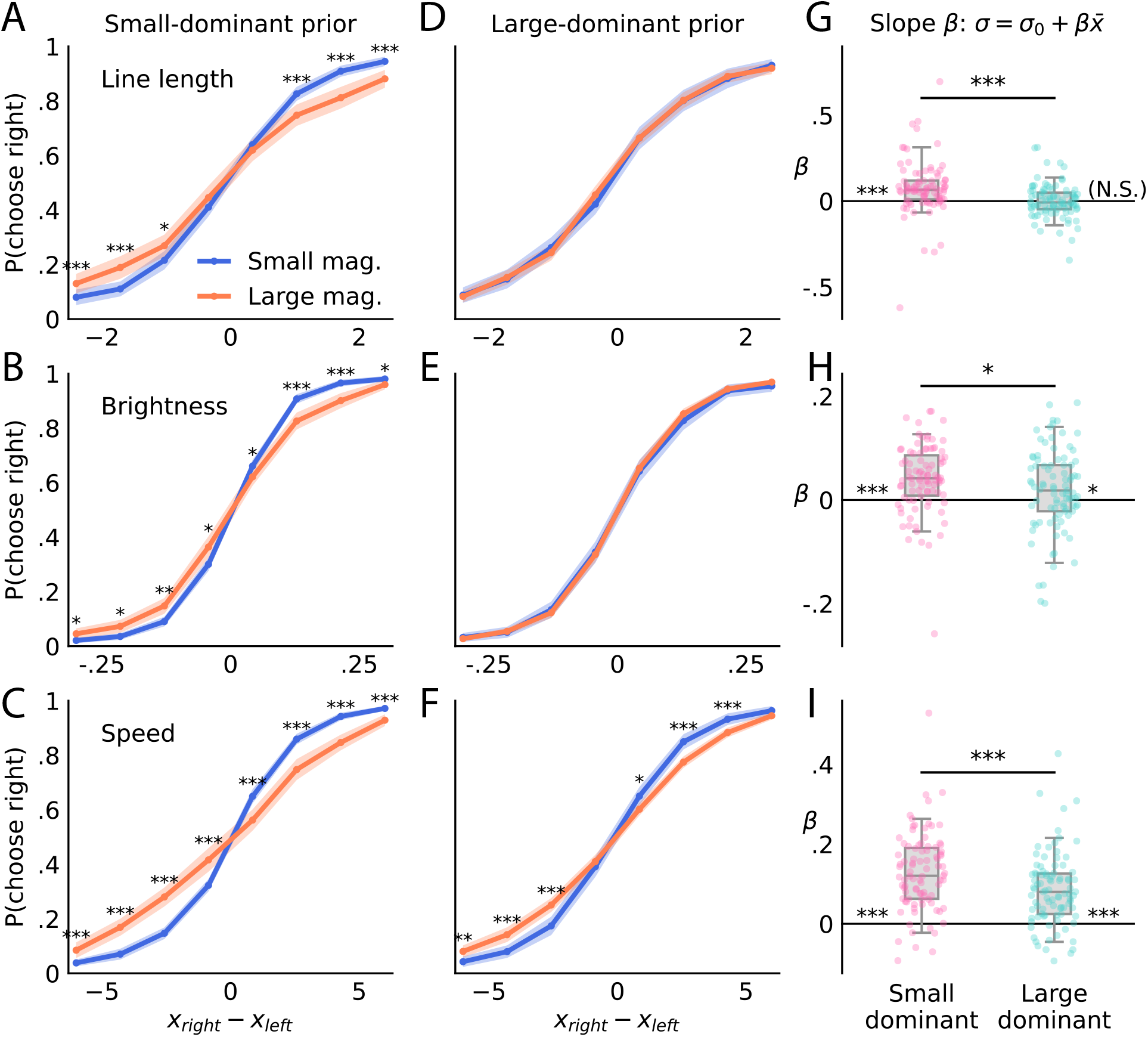
Suppression of Weber’s law when large magnitudes are more frequent. Proportion of ‘right’ choices as a function of the difference between left and right magnitudes, with the small-dominant **(A-C)** and the large-dominant prior **(D-F)**, with small (blue) and large (orange) magnitudes, in the line-length (A, D), brightness (B, E), and speed (C, F) experiments. The psychometric curves are sensitive to the prior and exhibit a suppression of Weber’s law with the large-dominant prior. Shaded areas show the 95% confidence interval. Abscissa units: differences in degrees of visual angles (dva; A, D), in brightness levels between 0 and 1 (B, E), and in dva/s (C, F). Stars indicate p-values of two-sided, paired Student’s *t*-tests corrected for multiple comparisons (Bonferroni-Holm-Šídák). We fit the psychometric curves to a logistic function with scale parameter 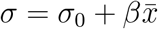 , where 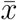 is the average magnitude in each trial. **(G-I)** show the best-fitting slopes *β* for the three experiments, in the two conditions. The slopes are significantly lower with the large-dominant prior, indicating a reduced sensitivity of the precision to stimulus magnitudes. Box plots indicate the median (line), quartiles (box), and 5th-95th percentiles (whiskers). Dots show individual participants. Stars near the abscissa indicate p-values of two-sided Student’s *t*-tests of equality with zero. ***: *P <* .001, **: *P <* .01, *: *P <* .05. N.S.: *P >* .05.

By contrast, with the large-dominant prior, no difference was detected between the psychometric curves of the small- and large-magnitude trials in the length and brightness experiments (Fig. 2D-E). Thus Weber’s law was suppressed, with these sensory modalities, when large magnitudes were more frequent than small magnitudes. In the speed experiment, participants remained more precise for small magnitudes, but by a lesser extent than with the small-dominant prior (Fig. 2F). To quantify these observations, we measured participants’ adherence to Weber’s law by fitting their responses to a logistic choice function whose scale parameter, *σ*, was allowed to linearly depend on the average magnitude at each trial. Specifically, 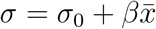 , where the slope parameter, *β*, determines the extent to which larger magnitudes are perceived with more imprecision than smaller magnitudes (we note that in this response model, *σ* is directly proportional to the JND; see Methods). For the three sensory modalities, in the small-dominant condition, the across-subjects average estimate of the slope *β* was significantly positive, consistent with Weber’s law (two-sided Student’s *t*-tests: length: *t*(87) = 4.4, *P* = 2.5 *×* 10^*−*5^; brightness: *t*(90) = 5.9, *P* = 6.5 *×* 10^*−*8^; speed: *t*(90) = 13, *P* = 1.7 *×* 10^*−*21^; Fig. 2G-I). Crucially, the slope *β* with the large-dominant prior was significantly lower than with the small-dominant prior, for all three sensory modalities (one-sided Welch’s *t*-tests: length: *t*(145.4) = 3.7, *P* = .0001; brightness: *t*(173.8) = 2.1, *P* = .017; speed: *t*(178.8) = 3.2, *P* = .0009; a Bayesian hierarchical model of statistical estimation supports these results; see Supplementary Information). In other words, manipulating the statistics of the sensory input modified the precision with which participants perceived the magnitudes of stimuli. Specifically, with the large-dominant prior their imprecision increased less with the magnitude than it did with the small-dominant prior.

In the length experiment, this resulted in slopes *β* which were not significantly different from zero (*t*(89) = 0.1, *P* = .921), and whose median was negative, i.e., the data suggest that a majority of subjects were more precise for large magnitudes than for small magnitudes, a behavior opposite to Weber’s law (Fig. 2G). In the brightness and speed experiments, the slope estimates remained positive on average (brightness: *t*(90) = 2.1, *P* = .04; speed: *t*(90) = 9, *P* = 4.2 *×* 10^*−*14^; Fig. 2H-I).

Two mechanisms could explain the softer slope *β* observed with the large-dominant prior: participants in this condition may have been more precise with large magnitudes, or less precise with small magnitudes. We found evidence for both mechanisms. With large magnitudes, the proportion of participants’ correct responses was significantly higher with the large-dominant than with the small-dominant prior, for all three sensory magnitudes (Fig. 3A; length: *t*(167.4) = 3.0, *P* = .002; brightness: *t*(142.5) = 2.8, *P* = .003; speed: *t*(172.1) = 2.4, *P* = .0099; one-sided Welch’s *t*-tests). Conversely, with small magnitudes, participants provided more correct responses when the prior was small-dominant, with a significant difference in the brightness experiment (brightness: *t*(154.6) = 3.5, *P* = .0003; length: *t*(176.0) = 1.4, *P* = .082; speed: *t*(174.3) = 1.4, *P* = .077). These results are consistent with a dynamic implementation of efficient coding, whereby the encoding precision of a stimulus is an increasing function of its contextual frequency.

**Fig. 3:**
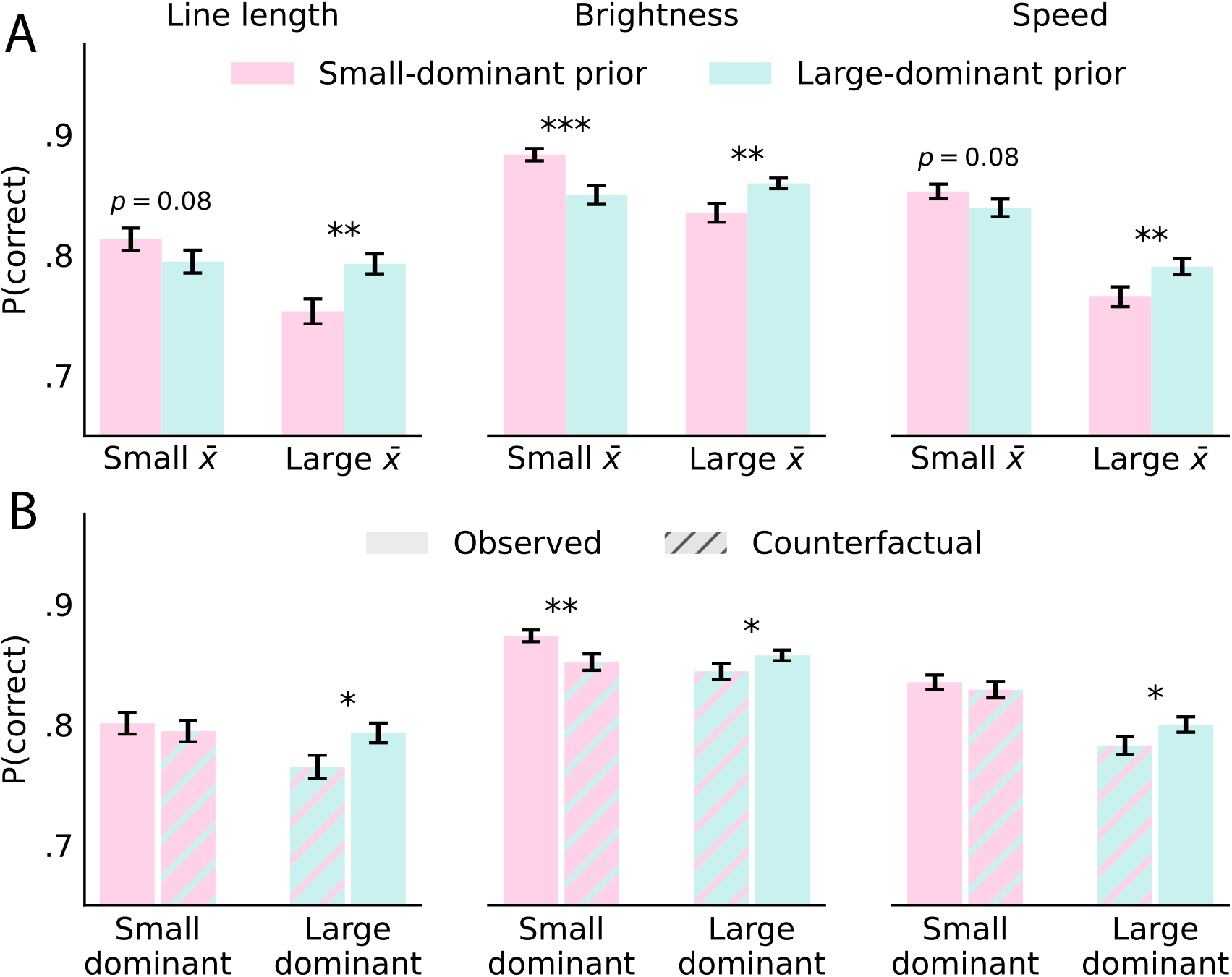
Participants adjusted their accuracies to the relative frequencies of small and large magnitudes, resulting in context-adapted performance. **(A)** Proportion of correct responses, for the three sensory modalities, with the small-dominant prior (pink) and the large-dominant prior (blue), with small magnitudes (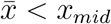 where 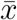 is the average magnitude in each trial and *x*_*mid*_ is the middle of the range), and with large magnitudes 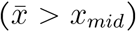. The accuracies for large (small) magnitudes are greater when large (small) magnitudes are more frequent, consistent with efficient coding. **(B)** Observed (solid-color bars) and counterfactual (hatched bars) overall proportion of correct responses in each condition. Counterfactuals are computed using the trials of one condition but the response statistics (illustrated in A) of the other condition. In each case, if participants adopted in one condition the accuracies for small and large magnitudes that they adopted in the other condition, then their overall performance would be inferior. Thus the participants’ context-sensitive accuracies enabled them to enhance their performance in the tasks. ***: *P <* .001, **: *P <* .01, *: *P <* .05.

We note that for a model participant who has perfectly learned the prior, efficient coding suggests that instead of a linear relation the noise scale parameter should take either one of two values, corresponding to small and large magnitudes (reflecting the prior shape; Fig. 1B). We found that this model provided a poorer fit than the linear model considered here. Its results were nonetheless consistent with our findings: in particular, the difference between the noise scales for large and small magnitudes was significantly smaller in the large-dominant condition. Moreover, in the large-dominant condition of both the line-length and brightness experiments, the two noise scales did not differ significantly (while they did in the small-dominant condition), indicating that the increase in imprecision at large magnitudes had vanished. This complementary analysis strengthens our conclusions (see Supplementary Information).

In short, the encoding of stimuli by participants appeared context-dependent. Did this adaptability enable them to increase their performance in the task? To measure this, we computed counterfactual response behavior by estimating the empirical accuracies in a given condition (separately for small and large magnitudes and for the tested magnitude differences, |*x*_*right*_ *− x*_*left*_|). We then applied these response statistics obtained in a given condition to the trials of the other condition. Informally, we thus estimated how well the ‘small-dominant behavior’ fared with the large-dominant prior, and conversely. We then compared this counterfactual performance to the actual, observed performance. In all cases the accuracies adopted by the participants in a given condition increased their performance in this condition, as compared to the counterfactual scenario in which they adopted the accuracies chosen in the other condition, with significant differences in most cases (Fig. 3B; permutation test, *N*_*perm*_ = 10^5^: small-dominant prior, lengths: *P* = .30, brightness: *P* = .0054, speed: *P* = .25; large-dominant prior, lengths: *P* = .014, brightness: *P* = .049, speed: *P* = .039). Therefore, participants’ behavior was consistent with a dynamic efficient coding mechanism whereby the differential precision of magnitude encoding was adapted across experimental conditions, in order to improve the number of correct responses in each condition—and thus, to increase the obtained reward in each experiment.

Finally, we surmised that this context-dependent encoding should also affect the temporal dynamics of choice behavior. With the small-dominant prior, we found that participants’ response times with small magnitudes were generally shorter than with large magnitudes (Fig. 4A-C). Hence trials with large magnitudes were characterized both by more erroneous choices and by slower responses. This is consistent with an evidence-accumulation model in which the drift rate is larger for small magnitudes than for large magnitudes, suggesting stronger sensory evidence for the more frequent magnitudes. The pattern of response times observed with the small-dominant prior was inverted with the large-dominant prior: indeed in this condition participants were faster at responding to large magnitudes than to small magnitudes (Fig. 4D-F). Overall this suggests that the strength of the sensory evidence was adapted, in each condition, to the current frequencies of large and small magnitudes.

**Fig. 4:**
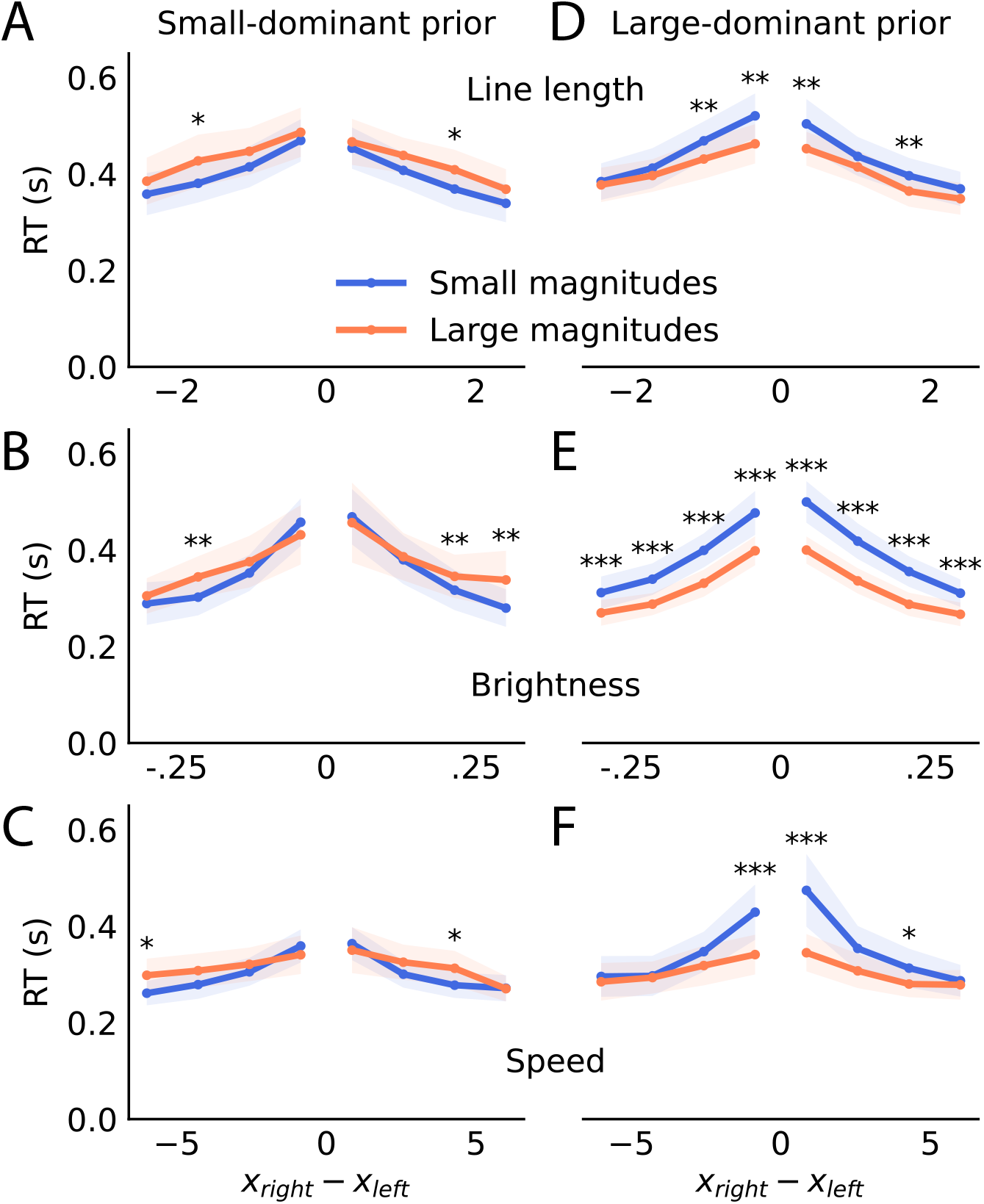
More frequent magnitudes elicit faster responses. Response times as a function of the difference between left and right magnitudes, with the small-dominant **(A-C)** and the large-dominant prior **(D-F)**, with small (blue) and large (orange) magnitudes, in the line-length (A, D), brightness (B, E), and speed (C, F) experiments. The magnitudes that dominate in the prior result in shorter response times. Shaded areas show the 95% confidence interval. Abscissa units as in Fig. 2. Stars indicate p-values of two-sided, paired Student’s *t*-tests corrected for multiple comparisons (Bonferroni-Holm-Šídák). ***: *P <* .001, **: *P <* .01, *: *P <* .05.

## Discussion

Our results show that one of the most robust regularities in psychophysics, Weber’s law, depends on the distribution of magnitudes encountered by observers. With three sensory modalities, when small magnitudes were more frequent, discrimination became less precise as magnitude increased, consistent with Weber’s law; making large magnitudes more frequent significantly reduced this dependence. The change in discrimination precision induced by the change of frequencies was not uniform across magnitudes: instead, accuracy increased in the region favored by the prior and tended to decrease in the other region. The accuracy profile obtained in one condition enabled a greater overall performance than the counterfactual accuracy profile transferred from the other condition. More frequent magnitudes also elicited faster responses. Together, our results show that recent stimulus statistics causally influence the relation between magnitude and discriminability; Weber’s law, therefore, is not an immutable property of sensory systems.

Furthermore, the observed pattern of changes in accuracies suggests a reallocation of limited representational resources toward more frequent magnitudes, increasing their precision at the expense of rarer magnitudes. This context-dependent redistribution of precision power was functionally adaptive (it improved task performance). Our results thus point to a dynamic implementation of the principle of efficient coding.

Most accounts of Weber’s law locate its origin in the internal encoding of external stimuli, for instance through logarithmic transformations, multiplicative noise, or specific evidence-accumulation mechanisms^3,8–12^. Our results are not inconsistent with these models (although they require them to be flexible enough to allow for the kind of adaptations that we exhibit). Regardless of the mechanisms mediating Weber’s law, our aim, rather, was to establish its rationale. In this regard our study is consistent with normative accounts that *derive* Weber’s law from the principle of efficient coding, combined with the observation that many sensory magnitudes exhibit a power-law distribution (i.e., skewed toward small magnitudes^6,22–26^). From this perspective, Weber’s law is the expected behavioral consequence of efficiently representing a particular class of environmental distributions. While these accounts infer this link theoretically, our study provides causal evidence that perceptual precision across magnitudes is shaped by stimulus statistics rather than immutably prescribed by Weber’s law.

Reframing Weber’s law in terms of efficient coding is fruitful because it is more expressive. The strictest formulation of Weber’s law posits a proportionality relation between the JND and the magnitude (*σ ∝ x*), but Fechner already noted that adding a constant was necessary for some sensory modalities, including brightness^1^; and further generalizations^2,3^ introduced an exponent, as *σ ∝* (*x* + *d*)^*α*^. This generalized form accounts for discrimination data with a broad range of sensory variables, in domains as diverse as gustatory, auditory, visual, vestibular, and numerosity perception^8,30–32^. Efficient coding immediately explains these results. Indeed, it predicts that the JND at a given magnitude should be proportional to a power of the magnitude’s probability^6^ (e.g., *σ ∝ f* (*x*)^*γ*^). Consequently, if stimulus magnitudes follow a generalized power-law distribution, the JND is predicted to be itself a generalized power law of the magnitude. For several of the sensory variables just mentioned, natural stimulus statistics have also been measured. Their empirical distributions are well described by generalized power laws, and the JND functions are indeed related to these distributions by a power transformation, as predicted by efficient coding^6,24,26^. Weber’s law thus emerges as a special case of this more general principle. Moreover, the distributions of some sensory variables, e.g., visual orientations, are not power laws, and the corresponding discrimination profiles do not obey Weber’s law; they are, however, consistent with efficient coding^6^.

Also readily explained by the efficient-coding account are the context-dependent accuracy profiles and the suppression of Weber’s law that we observe. But these suggest in addition that efficient coding can be implemented dynamically, rapidly adapting to changing stimulus statistics. This is consistent with behavioral studies showing that the range and shape of the stimulus distribution modulate the statistics of subjects’ responses^20,28,29,33–36^. This behavioral adaptability in discrimination tasks is presumably mediated by the flexibility of the underlying neural encoding of sensory inputs. Indeed neurophysiological studies have demonstrated the dynamic sensitivity of neural responses to stimulus statistics, for instance via gain rescaling in single sensory neurons^37–39^ and distributed range adaptation in sensory networks^27,40,41^. Some studies furthermore link these neural adaptations to corresponding behavioral changes^27,42^. In short, dynamic efficient coding in sensory networks enables adaptive behavior left unexplained by Weber’s law.

Providing further evidence for this reallocation, response times were shorter for the magnitudes that were more probable under the contextual prior. Analyses of response times in relation to Weber’s law are surprisingly scarce^11,12^. Pardo-Vazquez *et al*. account for response dynamics in discrimination tasks by requiring a power-law relationship between the stimulus magnitude and its internal representation (firing rates) in an evidence-accumulation model^12^. Our results suggests a different interpretation, whereby internal representations are tied to the probability distribution of stimuli, rather than to their magnitudes. In this view, power-law encoding is a consequence of power-law stimulus statistics rather than a fixed property of sensory systems.

## Methods

### Participants

The experiments were conducted remotely (online). We targeted 200 participants for each experiment (100 per condition; deviations from this target were due to the online recruitment platforms and out of our control; see Table S1 in Supplementary Information). Participants were restricted to above 18 years of age, located in the United States, and completed the experiment on desktop via web browser. See Table S1 for experiment-specific characteristics. Participants were compensated with a base compensation of $4.2 and a bonus based on performance (0.36¢ per correct response on learning and test trials). Hypotheses, design, and analyses were preregistered on AsPredicted prior to data collection (https://aspredicted.org/nv4j-wwp8.pdf, https://aspredicted.org/a3z26m.pdf, https://aspredicted.org/fy3ad7.pdf). The study protocol was approved by the Institutional Review Board (IRB) of Harvard University (protocol IRB15-2048). Informed consent was obtained from all participants.

As preregistered, participants were excluded if their accuracy fell below a threshold success rate (55% in the line length experiment, 60% in the brightness and speed experiments), and trials with RT >10s were excluded. In addition, participants whose estimates of the slope parameter *β* fell 5 SDs or more away from the average estimate were excluded from the analyses (see Supplementary Information for test results without these exclusions).

### Apparatus/Stimuli

All experiments were conducted online and developed in jsPsych 8.2.1^43^. The virtual-chinrest plugin^44^ was used to calibrate stimulus size according to viewing distance and monitor size/resolution.

#### Length

Stimuli in the length experiment were black horizontal lines of thickness 4 pixels and between 4.8-13.2 degrees of visual angle (dva) in length. The background color was white.

#### Brightness

Stimuli in the brightness experiment were circles 15dva in diameter with an annulus of width 0.375dva. Luminosity was specified on a linear scale spanning from 0 to 1, corresponding to black and white respectively. This was done to account for the nonlinear gamma encoding (sRGB) used by modern browsers. Values specified in this scale were transformed into 8-bit grayscale pixel values through a commonly used power-law approximation of the inverse sRGB transfer function^45,46^: *f* (*x*) = *x*^(1*/*2.2)^ * 255. The background was grayscale 2D Perlin noise with mean luminosity 0.5.

#### Speed

Stimuli in the speed experiment consisted of circular patches of horizontally drifting gratings with diameter 6dva. Gratings were broadband, spanning between 0.3 cycles/deg and 2.0 cycles/deg, with randomized phases and a power spectrum falling as *f* ^*−*2^. The drift speed of the gratings varied between 1-18 dva/s. The background color was uniformly #808080.

### Procedure

The experiments followed a two-alternative forced-choice (2AFC) procedure, consisting of 50 trials of training with feedback followed by 300 test trials without feedback. Stimuli were presented sequentially, one to the left and one to the right of the screen (with presentation order randomized; Fig. 1A). Participants were asked to fixate on a cross in the center of the screen and instructed to report if they thought the left or right stimulus was more intense (longer, brighter, or faster) on each trial using their keyboard. Additionally, participants were given untimed breaks every 100 trials and unlimited time to respond.

Stimulus magnitudes were generated as 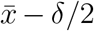 for the left stimulus and 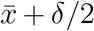 for the right, where 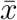 is the ‘base’ magnitude and *δ* is the difference in magnitude between left and right stimuli. *δ* values consisted of 8 evenly spaced values (see below for modality-specific values). 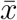 sampling was weighted such that in the *large-dominant* condition 20% of the 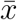 values were sampled uniformly from the lower half of the range (‘small’ range) and 80% uniformly from the upper half (‘large’ range), with the probabilities reversed in the *small-dominant* condition (Fig. 1B). Within each range, *δ*s were allocated to trials as equally as possible, and for each *δ* value the side appearing first was counterbalanced. Trial order was randomized. Each participant only saw a single experiment and condition.

#### Length

*δ* values consisted of 8 evenly spaced values between -2.4 and 2.4dva and 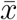 values were sampled from within 6-12dva. Stimuli were shown for 175ms with a 1000ms gap (fixation cross only) between stimuli. Lines were centered 6dva from the center of the screen randomly along a 90° arc (spanning either 315°-45° or 135°-225° depending on left/right placement).

#### Brightness

*δ* values consisted of 8 evenly spaced values between -0.3 and 0.3 and 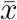 values were sampled from within 0.15-0.85. Stimuli were shown for 175ms with a 1000ms gap, and were offset horizontally 6.5dva from the center of the screen.

#### Speed

*δ* values consisted of 8 evenly spaced values between -6 and 6dva/s and 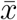 values were sampled from the largest range possible given the value of *δ* on that trial, such that all stimulus intensities fell within 1 and 18dva/s. Stimuli were shown for 350ms with a 500ms gap, and were offset horizontally 6dva from the center of the screen.

### Logistic choice model and JND

We fit, using maximum-likelihood estimation, a standard logistic choice function of the form *P* (responds ‘right’) = 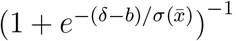 , where *δ* = *x*_*right*_ *− x*_*left*_ is the difference between the two magnitudes, and 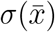 is an affine function of the average magnitude 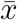 , as 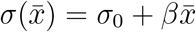. We note that *σ* is proportional to the just-noticeable difference (JND): indeed, defining the JND as the minimum difference *δ* needed to reach some accuracy *q* (e.g., *q* = 0.75), then in the absence of bias we have *q* = (1 + *e*^*−*JND*/σ*^)^*−*1^, i.e., 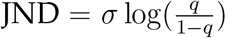 , that is, *σ* is proportional to the JND. Our analyses of participants’ adherence to Weber’s^1^l^*−*^a^*q*^w thus focus on the slope *β*, which measures how the JND linearly changes with stimulus magnitude. We also estimated the same model using a Bayesian hierarchical estimation method; this analysis supports our conclusions (see Supplementary Information).

