## Supplementary Information for "Efficient coding makes and breaks Weber’s law"

### Efficient coding makes and breaks Weber's law
#### Supplementary Information

Arthur Prat-Carrabin[1,⋆], Raphael Yamamoto[2], and Samuel J. Gershman[1]

[1]Department of Psychology and Center for Brain Science, Harvard University, Cambridge, MA
[2]Department of Computer Science, Haverford College, Haverford, PA

⋆**

August 10, 2026

### Efficient coding derivation and predictions

Here we present results pertaining to efficient-coding models, including how efficient coding relates to Weber's law. We consider an observer asked to estimate a magnitude $x$. The presentation of $x$ elicits in the brain of the observer a stochastic signal $r$ encoding $x$ (e.g., the vector of activities of a population of neurons). The distribution of $r$, $p(r|x)$, depends on $x$. The Fisher information is a statistical quantity that characterizes this conditional distribution, and provides a measure of the precision of the encoding, when the magnitude is $x$. It is defined as

$$I(x) = \mathbb{E}\left[\left(\frac{\mathrm{d}\ln p(r|x)}{\mathrm{d}x}\right)^2 \middle| x\right]. \quad (1)$$

A property of the Fisher information is that its inverse is the asymptotic variance of the Bayesian-mean estimator[1], which we denote by $\sigma^2(x)$. We note that this does not denote the variance of $x$, but the variance of the observer's estimate of $x$. The smaller this variance—i.e., the greater the Fisher information—the more precise the observer is about $x$.

We posit that the observer can choose the Fisher information, but with the additional assumption that a resource constraint prevents the observer from having unbounded information. Specifically, following the literature[2,3], we posit the constraint

$$\int I^{q/2}(x)\mathrm{d}x \leq K^{q/2}, \quad (2)$$

where $q > 0$. This constraint allows the Fisher information to be more precise about some magnitudes than about others, but under a general "budget" constraint on the allocation of precision.

The observer aims at minimizing a loss function, under this constraint. We choose the generalized loss function

$$\int \frac{f(x)^a}{I^{p/2}(x)} \mathrm{d}x, \tag{3}$$

where $f(x)$ is the prior over the magnitude, $p > 0$, and $a \in \{1, 2\}$. This loss function is a useful generalization that captures as special cases many objectives used in the literature, including the expected squared loss, and the mutual information[3]. The observer thus solves the following optimization problem:

$$\min_{I(x)} \int \frac{f(x)^a}{I^{p/2}(x)} \mathrm{d}x \quad \text{u.c.} \quad \int I^{q/2}(x) \mathrm{d}x \leq K^{q/2}. \tag{4}$$

Variational calculus provides the solution to this allocation problem. The solution is such that the Fisher information for the magnitude $x$ is proportional to a power of the relative frequency of $x$, as defined by the prior, i.e.,

$$I(x) \propto f(x)^{2\gamma}, \tag{5}$$

where $\gamma = \frac{a}{p+q}$. Therefore,

$$\sigma(x) \propto f(x)^{-\gamma}. \tag{6}$$

If the prior is a generalized power law, i.e., $f(x) \propto 1/(x + d)^\alpha$, we thus obtain

$$\sigma(x) \propto (x + d)^{\alpha\gamma}, \tag{7}$$

which yields Weber's law when $\alpha\gamma = 1$ and $d = 0$, and generalizations of the law otherwise.

### Participants

Table S1 summarizes participant recruitment, sample characteristics, exclusions, study duration, and compensation across the three experiments.

**Table S1: Participants characteristics across experiments.** Participants recruited through Amazon MTurk were CloudResearch[4] Approved Participants. F: female, M: male, NB: non-binary, O: other.

| | Line length | Brightness | Speed |
|---|---|---|---|
| N (*large-dom.* / *small-dom.*) | 199 (100 / 99) | 201 (101 / 100) | 195 (99 / 96) |
| N after exclusions | 184 (94 / 90) | 184 (92 / 92) | 183 (92 / 91) |
| Platform | Amazon MTurk | Amazon MTurk | CloudResearch Connect |
| Gender (F / M / NB / O) | 82 / 116 / 0 / 1 | 92 / 108 / 1 / 0 | 85 / 108 / 2 / 0 |
| Age, M (SD) | 44.80 (11.97) | 46.35 (12.69) | 42.55 (13.32) |
| Duration, median (IQR) | 22.84 (20.03-28.16) | 22.94 (20.19-27.73) | 22.30 (19.65-27.59) |
| Compensation, M (SD) | 5.58 (0.52) | 5.87 (0.51) | 5.21 (0.12) |

### Bayesian hierarchical estimation model

We estimate the logistic choice model with Bayesian statistical estimation methods. We use the following notations: $x_{si,right}$ and $x_{si,left}$ denote the right- and left-hand-side magnitudes shown to subject $s$ in trial $i$, $\bar{x}_{si}$ is the average magnitude in the trial (i.e., $\bar{x}_{si} = \frac{1}{2}(x_{si,right} + x_{si,left})$), $x_{min}$ and $x_{max}$ are the lowest and largest possible magnitudes in the experiment, $\sigma_{s,min}$ and $\sigma_{s,max}$ are two subject-specific noise-scale parameters (see below), $\mathrm{expit}$ is the sigmoid logistic function (i.e., $\mathrm{expit}(x) = \frac{1}{1+e^{-x}}$), $b_s$ is a subject-specific bias, and $d_{si}$ is the decision of subject $s$ in trial $i$ (where $d = 1$ means 'right' and $d = 0$ means 'left').

For numerical reasons, instead of computing the noise scale as $\sigma = \sigma_0 + \beta\bar{x}$ and fitting $\sigma_0$ and $\beta$, it is more convenient to define the parameters $\sigma_{s,min}$ and $\sigma_{s,max}$ as the values of the noise scale for the magnitudes $x_{min}$ and $x_{max}$, and interpolate in-between. In other words, instead of parameterizing the linear relationship with an intercept and a slope ($\sigma_0$ and $\beta$), we parameterize it with the values at the two extreme points ($\sigma_{s,min}$ and $\sigma_{s,max}$), which is strictly equivalent, but it enables us to enforce a positivity constraint by simply requiring $\sigma_{s,min} > 0$ and $\sigma_{s,max} > 0$.

Specifically, for each sensory modality, the model is defined by the following equations:

$$\begin{aligned} w_{si} &= \frac{\bar{x}_{si} - x_{min}}{x_{max} - x_{min}}, \\ \sigma_s(\bar{x}_{si}) &= (1 - w_{si})\sigma_{s,min} + w_{si}\sigma_{s,max} \\ p_{si} &= \mathrm{expit}\left(\frac{x_{si,right} - x_{si,left} - b_s}{\sigma_s(\bar{x}_{si})}\right), \\ \text{and } d_{si} &\sim \mathrm{Bernoulli}(p_{si}). \end{aligned} \tag{8}$$

**Hierarchical parameters** The parameters of the choice function include subject-level random effects. Specifically,

$$\begin{aligned} b_s &\sim N(b_0, \tau_b^2), \\ \log\sigma_{s,min} &\sim N(\log\sigma_{0,min,c}, \nu_{min,c}^2), \\ \log\sigma_{s,max} &\sim N(\log\sigma_{0,max,c}, \nu_{max,c}^2), \end{aligned} \tag{9}$$

where $c$ denotes the condition (small-dominant or large-dominant), and $b_0$, $\sigma_{0,min,c}$, and $\sigma_{0,max,c}$ capture group-level effects.

**Priors** Let $L = x_{max} - x_{min}$. The following priors were chosen:

$$\begin{aligned} b_0 &\sim N(0, L^2), \\ \tau_b &\sim N_+(0, L^2), \\ \sigma_{0,min,c} &\sim N_+(L/10, L), \\ \sigma_{0,max,c} &\sim N_+(L/10, L), \\ \nu_{min,c} &\sim N_+(0, 1), \\ \text{and } \nu_{max,c} &\sim N_+(0, 1), \end{aligned} \tag{10}$$

where $N_+$ denotes a normal distribution truncated to positive values.

**Slope** The condition-level slope of the noise scale was then recovered, as

$$\beta_{0,c} = \frac{\sigma_{0,max,c} - \sigma_{0,min,c}}{x_{max} - x_{min}}. \tag{11}$$

This slope captures the increase (or decrease) of the noise scale $\sigma$ with the magnitudes, for each condition $c$.

The statistical model was estimated using Stan with the HMC-NUTS sampler[5] (10 chains of 1,000 samples each, following 1,000 warmup iterations).

Figure S1A shows that for all three sensory modalities, the slope is larger in the small-dominant condition than in the large-dominant condition. Figure S1B shows the Bayesian posterior distributions for the difference in this slope across conditions, $\beta_{0,\text{small-dominant}} - \beta_{0,\text{large-dominant}}$. The posterior probability that this difference is negative is 0.05% in the line-length experiment, 0.86% in the brightness experiment, and 1.55% in the speed experiment.

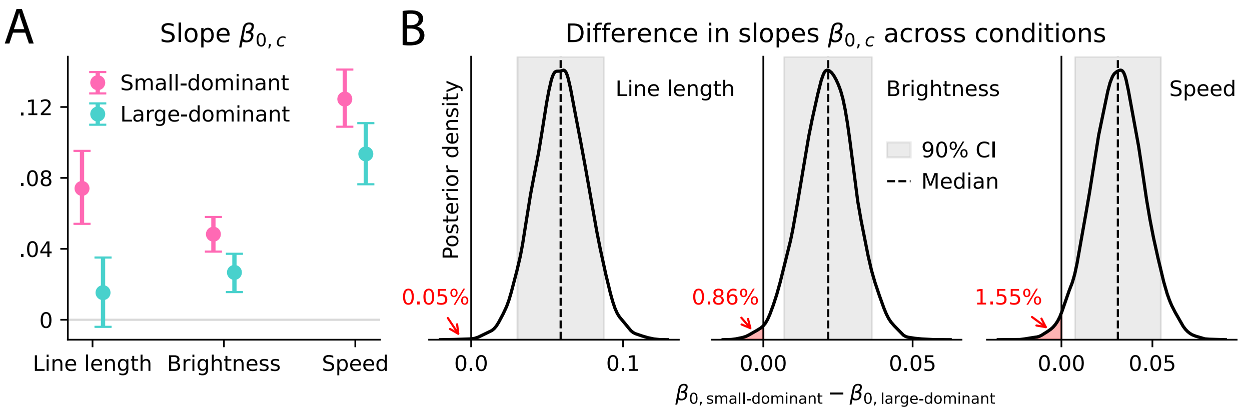


**Fig. S1: Hierarchical Bayesian estimates of the slope parameter $\beta$ in the linear relationship $\sigma = \sigma_0 + \beta\bar{x}$. (A)** Estimates (posterior medians and 90% credible intervals) of the slope parameter in each condition, $\beta_{0,c}$, with the three sensory modalities. **(B)** Bayesian posterior distribution of the difference between the slope parameters in the two conditions, $\beta_{0,\text{small-dominant}} - \beta_{0,\text{large-dominant}}$, in each experiment. Grey area: 90% credible interval. Dashed line: median. Positive values indicate that the subjects' noise scale increased faster with the magnitudes in the small-dominant condition than in the large-dominant condition. Red areas and percentages show the Bayesian probability of negative values.

### Tests

Table S2 shows the results of the preregistered, one-sided, two-sample $t$-tests of whether the best-fitting estimates of the slope parameter $\beta$ is lower in the small-dominant condition than in the large-dominant condition, when excluding or including outliers (estimates of $\beta$ five SDs away from the mean). The last column shows the Bayesian posterior probability that the difference in slopes, $\beta_{0,\text{small-dominant}} - \beta_{0,\text{large-dominant}}$, is negative, as estimated with the hierarchical model presented above (outliers were not excluded for this analysis). This Bayesian probability is also reported in Fig. S1B.

These results substantiate our conclusions that increasing the frequencies of large magnitudes and decreasing the frequencies of small magnitudes significantly impacted the discriminability of stimuli: specifically, in the large-dominant condition the scale of the imprecision grew less steeply with the magnitude.

**Table S2: Tests of whether the slope in the large-dominant condition exceeds the slope in the small-dominant condition.**

| | Excluding outliers | | Including outliers | | Bayesian probability |
|---|---|---|---|---|---|
| Modality | Statistic | $p$-value | Statistic | $p$-value | (incl. outliers) |
| Line length | $t(145.4) = 3.7$ | 0.00014 | $t(181.5) = 1.2$ | 0.122 | 0.05% |
| Brightness | $t(173.8) = 2.1$ | 0.017 | $t(172.3) = 1.9$ | 0.029 | 0.86% |
| Speed | $t(178.8) = 3.2$ | 0.00087 | $t(153.5) = 1.6$ | 0.054 | 1.55% |

### Alternative efficient-coding logistic model

Here we consider a logistic choice model in which the noise scale parameter, $\sigma$, is not a linear function of the magnitude. Instead, it takes one of two values: $\sigma_{small}$ in small-magnitude trials, and $\sigma_{large}$ in large-magnitude trials. We note that this is consistent with the efficient-coding prediction that the precision should be a power transformation of the prior (Eq. 6), in the case of a prior that is uniform over each half of the magnitude range, as in our experiments. Overall, the BIC of this 'two-noise-scales' model is higher by 226 points than the linear model; thus in the main text we report the results of the linear model.

Although it does not yield the best fit, we can nevertheless ask whether the best-fitting parameters obtained with this model are consistent with our conclusions. We thus look at the difference, $\sigma_{large} - \sigma_{small}$, between the noise scale parameters for the large- and small-magnitude trials; this difference is conceptually similar to the slope parameter $\beta$ of the linear model. For all three sensory modalities, we find that in the small-dominant condition, this quantity is significantly positive, i.e., the subjects are more imprecise for large magnitudes than for small magnitudes ($\sigma_{large} > \sigma_{small}$), consistent with Weber's law (Fig. S2).

This difference between the noise in large vs. small magnitudes, however, is significantly smaller in the large-dominant condition, for the three sensory modalities. As a result, in the line-length and brightness experiments, the difference itself ($\sigma_{large} - \sigma_{small}$) is not significantly different from zero (Fig. S2).

Overall, these results are consistent with those obtained with the linear model, and they strengthen our conclusions: while in the small-dominant condition participants were more precise for small magnitudes than for large magnitudes, this difference was suppressed in the large-dominant condition. In other words, manipulating the relative frequencies of magnitudes changed the precision with which participants were able to discriminate the magnitudes, consistent with a dynamic implementation of efficient coding.

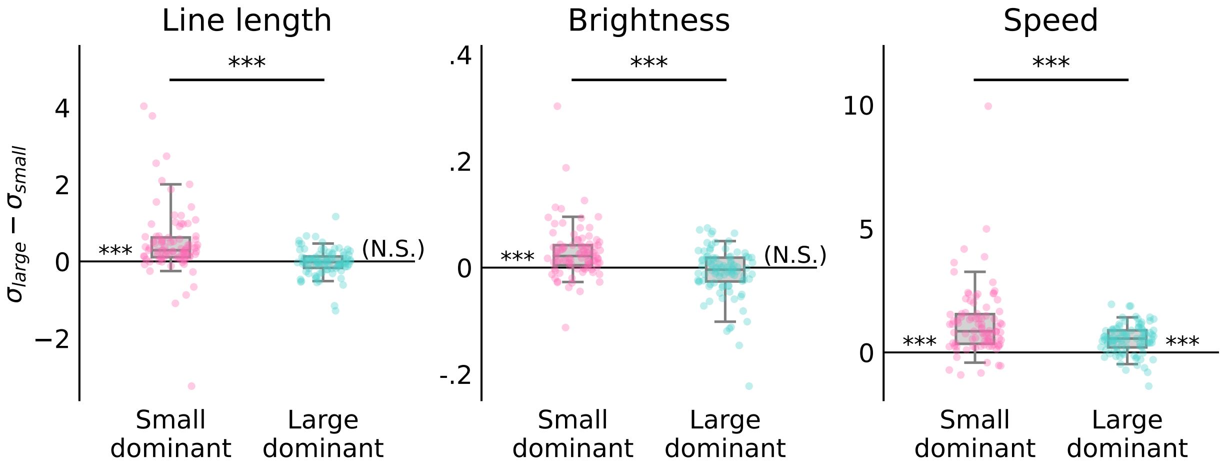


**Fig. S2: Suppression of the difference between noise levels for large and small magnitudes, in a choice model with two discrete noise levels.** Difference between the two noise scale parameters, $\sigma_{large} - \sigma_{small}$, in the small-dominant and large-dominant conditions, for the three experiments. Box plots indicate the median (line), quartiles (box), and 5th-95th percentiles (whiskers). Dots show individual participants. Stars near the abscissa: p-values of two-sided Student's $t$-tests of equality with zero; stars across box plots: p-values of one-sided Welch's $t$-tests of equality across conditions. ***: $P < .001$, **: $P < .01$, *: $P < .05$. N.S.: $P > .05$. Compare to Fig. 2G-I.
